# Allele-specific splicing modulates protein isoforms and Alzheimer’s risk

**DOI:** 10.64898/2026.02.13.705796

**Authors:** Alison J. King, Kofi Amoah, Laixing Zhang, Jae Hoon Bahn, Ryan M. Barney, Giovanni Quinones-Valdez, Xinshu Xiao

**Affiliations:** Bioinformatics Interdepartmental Program, University of California, Los Angeles, USA; Department of Integrative Biology and Physiology, University of California, Los Angeles, USA

## Abstract

Despite growing catalogs of genetic variation linked to human traits and diseases, the functional impact of most genetic variants remains poorly understood. Alternative splicing, particularly in the human brain, represents a key layer of post-transcriptional regulation that may mediate genetic effects on gene expression and protein diversity. In this study, we systematically map allele-specific alternative splicing (ASAS) events in postmortem brain tissues from the Mount Sinai Brain Bank cohort, identifying hundreds of genetically regulated splicing events across four brain regions. Using a concordance-based method, we nominate over 500 putative functional SNPs associated with ASAS, many of which overlap splicing QTLs (sQTLs), RNA-binding protein binding sites, and GWAS loci for Alzheimer’s disease (AD), brain traits, and immune phenotypes. ASAS events are enriched in genes involved in mitochondrial function and frequently occur in 5′ untranslated regions (5′ UTRs), where they are associated with protein quantitative trait loci (pQTLs), alternative start codons, and isoform-specific domain changes - highlighting an underappreciated mechanism through which noncoding variants can influence translation and proteome complexity. Importantly, we also identify a subset of ASAS events exhibiting disease-specific splicing patterns in AD brains, including functional SNPs with opposing splicing effects between AD and control groups in genes implicated in mitochondrial function and neuronal signaling. Together, our results provide a brain-specific, splicing-resolved map of regulatory variation and uncover novel mechanisms linking genetic variation to transcript and protein-level changes in AD. This work highlights the importance of allele-specific splicing analysis for interpreting noncoding variation in complex human disorders.

## Introduction

Large-scale datasets have enabled genome-wide association studies (GWAS) to identify thousands of genetic variants potentially contributing to complex traits and diseases (Buniello et al. 2019). However, the vast majority of GWAS-identified variants have no known function, and linkage disequilibrium (LD) structure often makes it difficult to determine which variants are truly causal. Genetic variants may influence traits through a variety of regulatory mechanisms, including effects on gene expression, RNA splicing, or protein abundance. These complexities make it challenging to infer functional mechanisms solely from GWAS data. Thus, elucidating how specific variants exert their effects is essential for uncovering the biological basis of complex traits and diseases.

One biological process frequently affected by genetic variants is alternative splicing, a post-transcriptional mechanism that enables a single gene to give rise to multiple transcript isoforms through differential inclusion of exons and introns. Over 90% of human genes undergo alternative splicing, contributing significantly to phenotypic and functional diversity (Wang et al. 2008; Pan et al. 2008). This process is tightly regulated and highly tissue-specific, with the brain exhibiting particularly complex splicing programs (Yeo et al. 2004). Splicing regulation depends on a coordinated interplay between *cis*-regulatory elements, such as splice sites, enhancers, and silencers, and *trans*-acting proteins, primarily RNA-binding proteins (RBPs). Consequently, genetic variants that disrupt *cis*-elements can have significant effects on splicing decisions, with potential downstream consequences for gene function.

Alternative splicing has been implicated in numerous human diseases (Li et al. 2016; Scotti and Swanson 2016). In Alzheimer’s Disease (AD), transcriptome analyses have identified widespread changes in splicing patterns (Raj et al. 2018), including dysregulated splicing events in multiple AD-associated genes (Raj et al. 2018; Course et al. 2023; Malik et al. 2013; Goedert et al. 1989). Moreover, mitochondrial dysfunction (Fairbrother-Browne et al. 2021; D’alessandro et al. 2025) and aberrant immune activation (Gate et al. 2020; Prater et al. 2023), both molecular hallmarks of AD, have been linked to splicing regulation. Prior work has identified genetically regulated splicing events in genes involved in these pathways (Boussaad et al. 2020; Rotival et al. 2019), raising the possibility that splicing dysregulation may contribute causally to AD pathogenesis. However, the extent to which genetic variants drive such splicing changes, and how they relate to disease-relevant molecular phenotypes, remains incompletely understood.

One powerful strategy to identify *cis*-regulatory effects on splicing is to examine allele-specific alternative splicing (ASAS), where one allele of a heterozygous single nucleotide polymorphism (SNP) is preferentially associated with a specific splicing pattern. ASAS enables the detection of regulatory variants by comparing allelic effects within the same individual, controlling for environmental and *trans*-acting influences. While previous studies have mapped genome-wide ASAS patterns in human tissues (Amoah et al. 2021), few have examined allele-specific splicing in the context of neurodegenerative disease. Another common approach is the mapping of splicing quantitative trait loci (sQTL), but these analyses typically require a large amount of data to detect statistical associations. Deep learning models (Jaganathan et al. 2019) have also been applied to predict splicing effects, but these are often less interpretable with limited performance in pinpointing splicing-altering SNPs.

In this study, we leverage RNA-seq and genotype data from the Mount Sinai Brain Bank (Wang et al. 2018) (MSBB) to identify and characterize genetically regulated ASAS events in postmortem human brain tissues. Our goal was to investigate how genetic variation influences splicing regulation in the brain and to explore the potential role of these splicing events in neurodegeneration. Our findings reveal widespread genetic modulation of splicing in the brain, particularly within the 5’ UTR, underscoring ASAS as an important mechanism linking genetic variation to AD genes, protein abundance, and isoform diversity.

## Results

### Landscape of ASAS events in brain regions

We previously developed a method (Li et al. 2012) to identify ASAS events within one RNA-seq dataset by integrating analyses at both single-nucleotide and transcript-isoform levels (Methods). Briefly, this approach compares allelic biases of heterozygous SNPs located in alternatively spliced regions against those in constitutively spliced regions within the same gene. ASAS events are pinpointed when SNPs within alternatively spliced regions exhibit significant allele-specific bias in read counts relative to those in constitutive regions (Methods). ASAS reflects the existence of genetic regulation on splicing of the associated exons. It should be noted that the SNP showcasing the allele-specific bias, i.e., the tag SNP, may or may not be the functional SNP that causes the ASAS pattern. We applied this method to RNA-seq data collected from the Mount Sinai Brain Bank (Wang et al. 2018) (MSBB). Following quality controls (Methods), we analyzed a total of 1,047 samples, originating from 4 brain regions and 293 individuals. In total, we identified 678 unique tag SNPs associated with 442 ASAS exons (Supplemental Table S1), with some exons linked to multiple tag SNPs. About 20% of these SNPs and 24% ASAS exons overlapped those identified in our previous work within tissues from the Genotype-Tissue Expression (GTEx) project (Amoah et al. 2021). This finding validates our results against established datasets while revealing a distinct set of splicing events unique to the aging and AD brain context.

The distribution of the number of ASAS events per sample was largely consistent across brain regions (Figure 1A). In addition, when assessing the overlap of unique ASAS events across regions, we observed that the largest set comprised events shared by all four brain regions, indicating substantial consistency in ASAS across these regions (Figure 1B). This trend still holds when restricting the comparison to ASAS events testable in all regions (Supplemental Fig. S1). To assess the relative contributions of inter-individual, inter-regional, and *APOE* genotype differences to ASAS variation, we implemented a linear mixed model as previously described (Amoah et al. 2021) (Methods). The model revealed that individual-level differences accounted for more variance in allelic bias at tag SNPs than differences between brain regions or *APOE* genotype, with the latter two accounting for similar proportions (Supplemental Fig. S2). This pattern likely reflects the substantial genetic heterogeneity across the cohort, contrasting with the relatively high transcriptomic similarity shared among the sampled brain regions. Notably, for a small subset of tag SNPs, *APOE* genotype emerged as the dominant source of variation. These SNPs map to genes with established AD relevance or *APOE* interactions, such as *VSTM2A* and *SNHG14* (Arzouni et al. 2020; Fitz et al. 2021), as well as the mitochondrial metabolic regulator *ACADM*.

**Figure 1.**
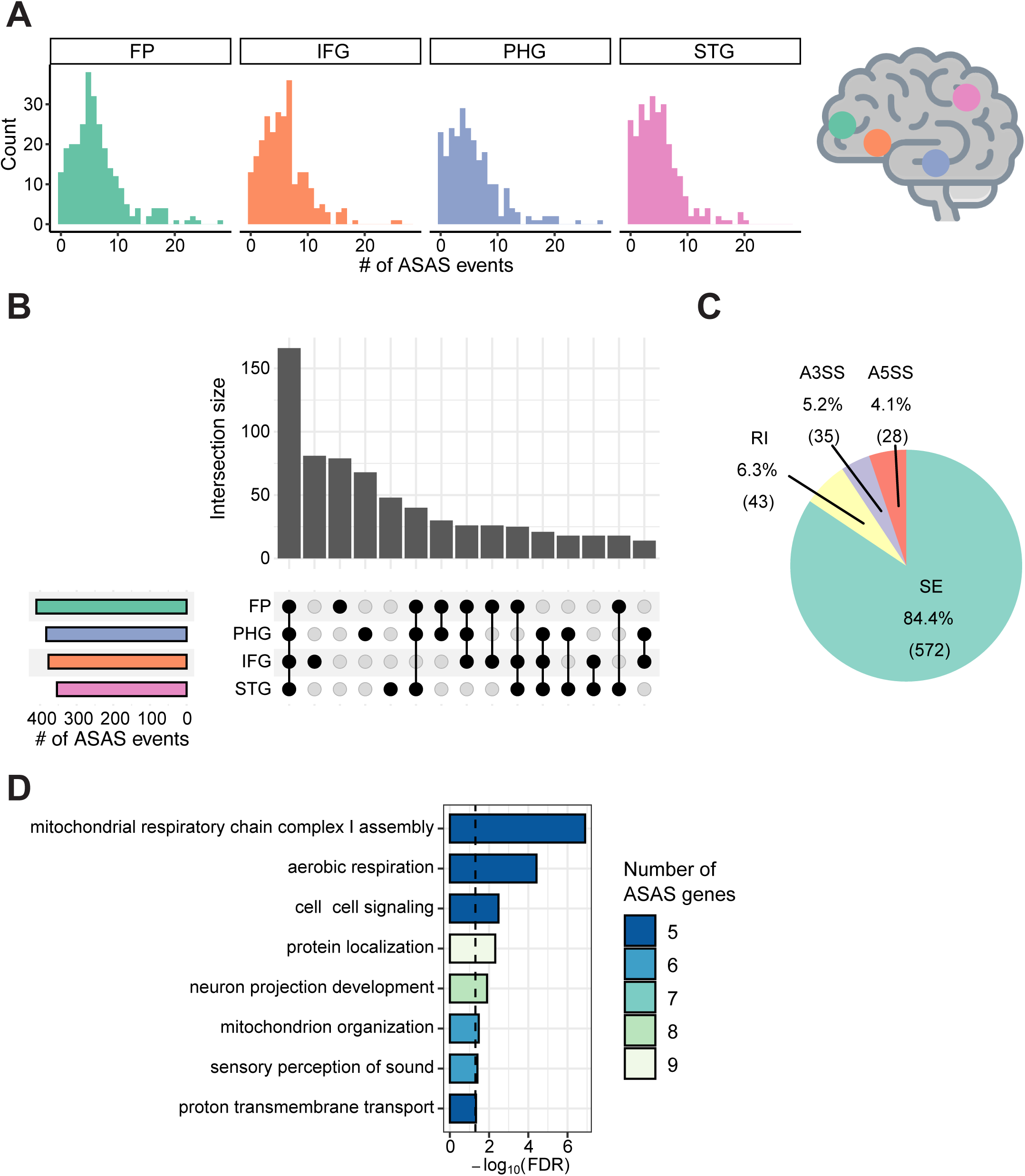
Landscape of ASAS events in brain regions. (A) Number of ASAS events per sample, separated by brain region: STG: superior temporal gyrus; PHG: parahippocampal gyrus; IFG: inferior frontal gyrus; FP: frontal pole. (B) Overlap of ASAS events across brain regions. (C) Types of alternative splicing for all unique ASAS events. SE: skipped exon; RI: retained intron; A3SS: alternative 3’ splice site; A5SS: alternative 5’ splice site. (D) GO terms enriched among genes with ASAS events, colored by the number of genes associated with each GO term. 10,000 sets of random background genes with similar expression level and gene length were used to generate a null distribution. The p-value was calculated based on the null distribution, with an FDR threshold of 0.05 (Methods).

Among the ASAS events, the vast majority were classified as skipped exons (SE), which accounted for 84.4% of all events (Figure 1C). The remaining events consisted of alternative 5’ splice sites (A5SS), alternative 3’ splice sites (A3SS), and retained introns (RI), each contributing a similar proportion, consistent with previous observation (Amoah et al. 2021). To gain insight into the biological functions impacted by ASAS, we performed Gene Ontology (GO) enrichment on genes harboring ASAS events (Methods). At an FDR threshold of 0.05 (used for all ensuing statistical tests), enriched GO terms highlighted several biological processes relevant to brain function and disease, such as mitochondrial respiratory chain complex I assembly, mitochondrion organization, proton transmembrane transport, aerobic respiration, neuron projection development, and cell-cell signaling (Figure 1D). Given the mitochondrial-related enrichment, we compared genes harboring ASAS events to a curated set of genes encoding proteins in the mitochondrial proteome (Rath et al. 2021). Using background genes that were testable for ASAS but not found to be significant, we observed a significant enrichment of ASAS genes among those with mitochondrial localization (hypergeometric test, p = 0.0267). This enrichment of mitochondrial-related terms reinforces prior evidence that mitochondrial dysfunction plays a central role in aging and AD (Fairbrother-Browne et al. 2021; D’alessandro et al. 2025; Holper et al. 2019). In particular, dysregulation of the mitochondrial chain complex I has been observed in AD (Rhein et al. 2009; Aksenov et al. 1999) and has been explored as a therapeutic target in preclinical AD models (Stojakovic et al. 2021).

### Identification of putative functional SNPs regulating ASAS events

To identify candidate functional SNPs regulating ASAS events, we applied a concordance-based method developed in our previous study (Amoah et al. 2021) (Methods). This method prioritizes for functional SNPs by evaluating the concordance between SNP genotypes and allele-specific splicing patterns across individuals. Specifically, for each candidate SNP located within the same gene as an ASAS-associated tag SNP, a concordance score is calculated. This score quantifies how well the genotype of the candidate SNP aligns with the expected direction and magnitude of allelic bias observed at the tag SNP. The key assumption is that a functional SNP, when heterozygous, will consistently produce a predictable allele-specific splicing pattern across the cohort. We used this framework in two analytical modes: (1) across all samples within a brain region to capture broadly acting variants, and (2) with AD-stratified groups to account for disease-specific effects. For the latter, we categorized individuals based on their CERAD neuropathology scores (Morris et al. 1989). Those classified as “normal” were considered controls, and those with “definite AD” were classified as AD cases. After quality control (Methods), this analysis included over 60 controls and 85 AD samples per brain region (Supplemental Fig. S3). Detailed cohort demographics, including post-mortem interval, age, sex, and *APOE* genotype, are provided in Supplemental Fig. S3. As expected, the frequency of the *APOE* ε4 allele, the strongest genetic risk factor for late-onset AD, was significantly higher in cases compared to controls (odds ratio = 6.52, p < 2.2 x 10^-16^).

We identified 501 putative functional SNPs across all regions and conditions, corresponding to 109 ASAS exons at an FDR threshold of ≤ 0.05 (Supplemental Table S2). Because these SNPs are predicted to modulate splicing in *cis*, we hypothesized that they would be located in closer proximity to their associated ASAS exons, consistent with prior studies showing that splicing regulatory elements often reside near their target exons (The GTEx Consortium 2020; Zhang et al. 2020). Indeed, functional SNPs exhibited a strong positional bias, with the majority located within a few hundred base pairs of the associated ASAS exon (Figure 2A). To benchmark this observation, we generated a null distribution by randomly sampling ‘testable’ SNPs (100 iterations). Unlike the functional set, these random samples exhibited no positional enrichment near splice sites, confirming the specificity of the observed bias. This spatial enrichment provides additional support for the regulatory relevance of the predicted functional SNPs.

**Figure 2.**
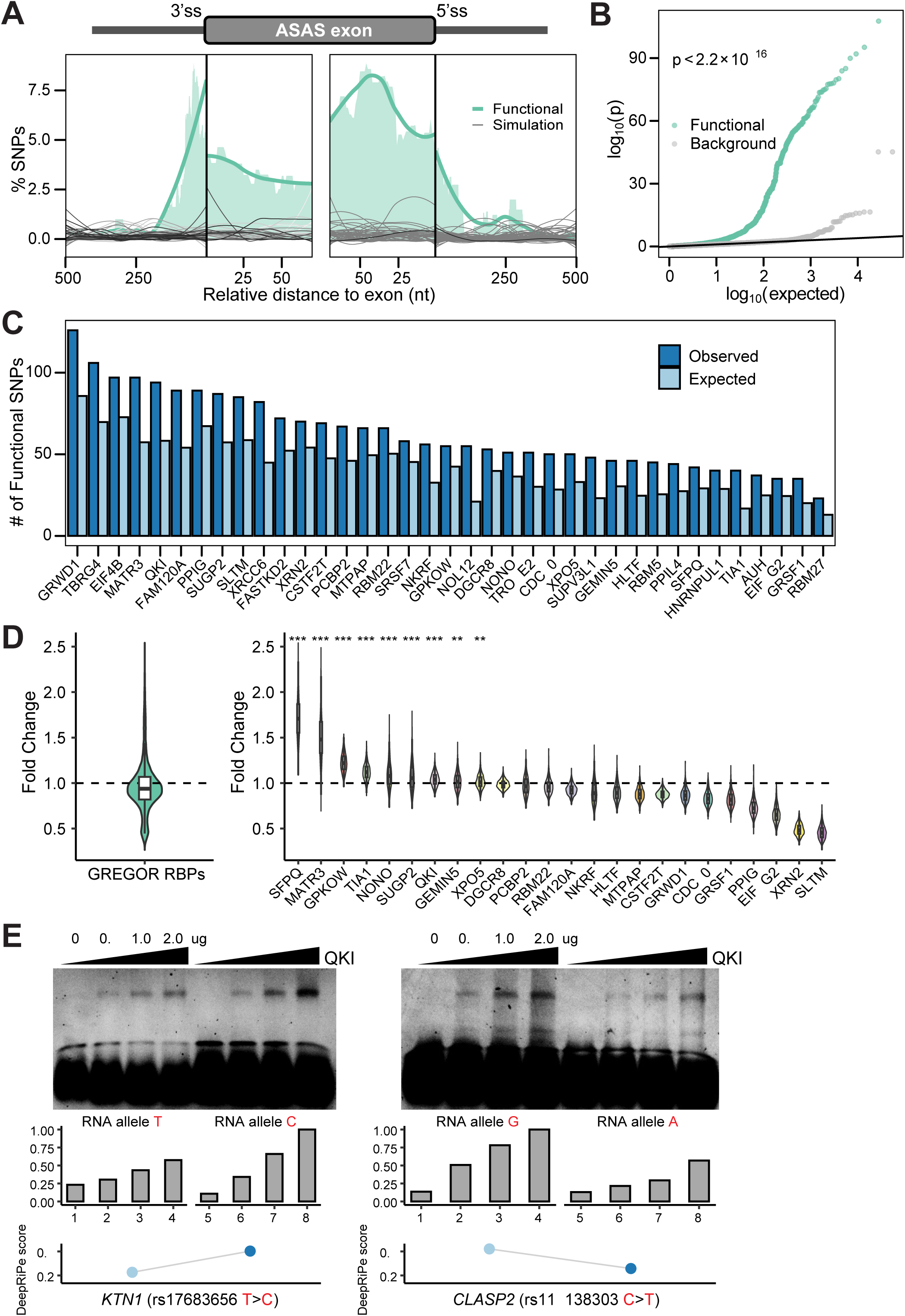
Identification of putative functional SNPs regulating ASAS events. (A) Distances of putative functional SNPs to their associated ASAS exon boundaries normalized by the total number of testable SNPs at each position. Green curve is the smoothed trend line of the shaded area representing the SNP density. Gray curves correspond to 100 sets of randomly sampled SNPs (Methods). (B) Quantile-quantile (Q-Q) plot of p-values for putative functional SNPs overlapping all tested brain GTEx sQTLs. A matched background distribution was generated by randomly sampling the same number of SNPs from all tested SNPs (Methods). (C) RBPs with significantly enriched binding sites (expected versus observed) among functional SNPs and their LD proxies, evaluated using GREGOR (Genomic Regulatory Elements and Gwas Overlap algoRithm) (Methods). (D) Distribution of predicted fold changes in RBP binding affinity scores between reference and alternative alleles of functional SNPs. Fold changes are shown for all significantly enriched RBPs combined (left) and for each individual RBP (right) identified in (C) with a DeepRiPe model. Fold change was calculated as the median score change among functional SNPs divided by the median from 1,000 random SNP sets (Methods), evaluated with a one-sided Mann-Whitney U test. ***p < 2.2 x 10^-16^. **p < 0.01. (E) EMSA validating allele-specific binding of QKI to ASAS-associated SNPs. The amount of QKI protein used is shown above the gel. The DeepRiPe predictions are shown below each allele.

To further substantiate these findings, we assessed whether our predicted functional SNPs were captured as significant sQTLs in brain tissues from the GTEx project (Methods). Our analysis revealed that the predicted functional SNPs were significantly enriched for low p-values in GTEx sQTLs compared to a background of all tested variant-gene pairs (Figure 2B). This observation provides orthogonal support for the potential regulatory roles of the SNPs identified by our method.

Splicing regulation is often governed by the interaction of RBPs with pre-mRNA and disruption of these interactions by genetic variants can alter splicing patterns (Van Nostrand et al. 2020a). To assess whether our predicted functional SNPs are enriched within RBP-binding sites, we intersected them with eCLIP-defined RBP-binding peaks and assessed enrichment using the GREGOR algorithm (Schmidt et al. 2015) (Methods). GREGOR compares the observed overlap to that of matched control SNPs, accounting for confounding factors such as the number of variants in LD, minor allele frequency, and distance to nearest gene. This analysis revealed significant enrichment for 37 RBPs (Figure 2C), many of which have established roles in RNA processing and splicing regulation (Van Nostrand et al. 2020b; Boyle et al. 2023; Coelho et al. 2015). For example, MATR3, one of the top enriched RBPs, has been shown to be involved in splicing and neurodegenerative diseases (Coelho et al. 2015; Khan et al. 2024). We also observed significant enrichment for QKI, which is essential for myelination and reported to be upregulated in AD brains (Farnsworth et al. 2016), and TIA1, a stress-granule component that interacts directly with Tau pathology (Ash et al. 2021). This enrichment suggests a specific mechanism where functional SNPs disrupt the core recognition motifs for these neuro-regulatory factors. Consequently, the resulting allele-specific loss of RBP binding does not just alter splicing, but may directly contribute to the myelin defects and proteostatic stress characteristic of the AD brain.

To determine whether these predicted functional SNPs modulate splicing by altering RBP binding, we used DeepRiPe (Ghanbari and Ohler 2020) to predict the effect of each SNP on RBP binding scores. We then compared these predicted changes to matched control variants (Methods). A number of RBPs with enriched binding sites had significant shifts in binding scores at functional SNPs (Figure 2D), supporting a model in which these variants may mediate allele-specific splicing by perturbing RBP-RNA interactions. To directly assess RNA-RBP interactions, we performed electrophoretic mobility shift assays (EMSA) using recombinant QKI protein and RNA probes representing its putative target SNPs. For each SNP, we synthesized RNA oligonucleotides corresponding to the reference and variant alleles. As shown in Fig. 2E, QKI exhibited dose-dependent binding to the RNA probes. Consistent with the DeepRiPe predictions, the variant alleles significantly altered QKI binding affinity relative to the references. These results provide direct biochemical evidence that these SNPs functionally alter protein-RNA interactions, supporting our hypothesis that genetic variants can drive splicing changes by altering RBP recognition motifs.

### ASAS enrichment in AD-relevant genes and GWAS risk loci

To investigate the potential disease relevance of ASAS events, we first asked whether genes harboring ASAS events overlapped with known AD-relevant gene sets. We observed significant enrichment of ASAS genes among both candidate therapeutic targets and predicted AD-relevant genes (Britton et al. 2023) (Figure 3A), suggesting that ASAS may affect splicing in genes implicated in AD pathogenesis. We then examined whether ASAS-associated SNPs (including both tag SNPs and predicted functional SNPs) were enriched in GWAS loci. Direct overlap revealed over 40 ASAS-associated SNPs were also reported as lead GWAS variants, a proportion significantly higher than expected by chance (Figure 3B, Methods). These results indicate that ASAS-associated SNPs likely intersect with genetic risk loci for human diseases.

**Figure 3.**
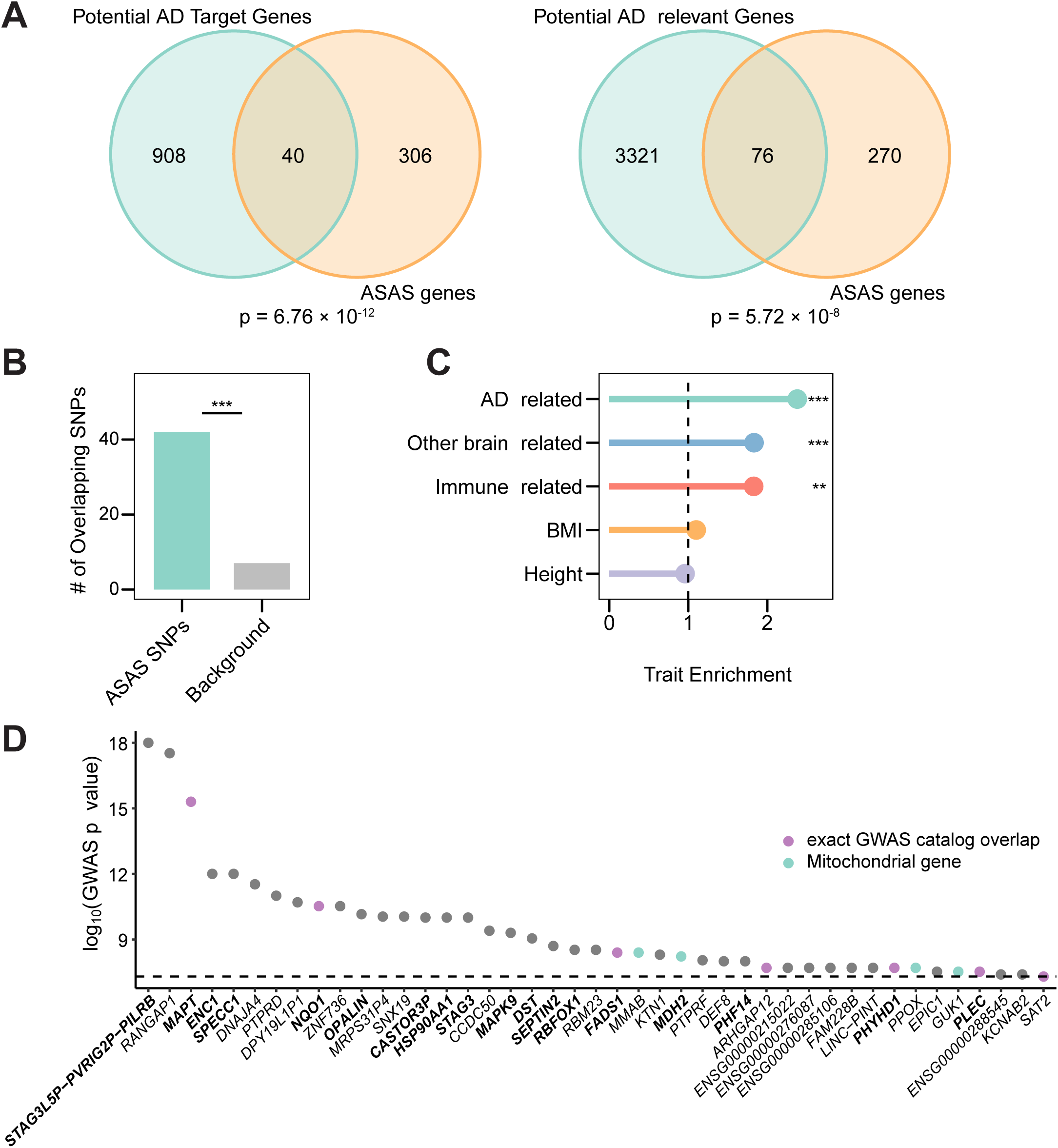
ASAS is enriched in AD-relevant genes and GWAS loci. (A) Overlap of genes harboring ASAS events with nominated AD targets (left) or potential AD-relevant genes (right). The significance was evaluated with a hypergeometric test compared to a background of annotated genes. (B) Number of ASAS-associated SNPs that overlap GWAS SNPs. The background set contains the same number of random common SNPs from genes without ASAS events. ***p = 2.76 x 10^-7^ (Fisher’s exact test). (C) Enrichment of ASAS-associated SNPs in LD with at least one GWAS SNP and within a 200 kb window, analyzed for GWAS category groups of related GWAS traits (Methods). The enrichment was calculated as the proportion of ASAS-associated SNPs in LD with the GWAS SNPs compared to that of the same number of randomly sampled common SNPs, evaluated with Fisher’s exact test. ***p < 0.001; **p < 0.01. (D) Genes harboring ASAS-associated SNPs in LD with GWAS SNPs from the AD-related trait group in (C); Y-axis shows the most significant p-value from the associated GWAS SNPs. AD-relevant genes are shown in bold, mitochondrial-related genes are shown in green, and genes with ASAS-associated SNPs exactly overlapping a GWAS SNP are shown in purple. Ensembl IDs are shown for genes without approved HGNC symbols.

Although several methods have been developed to estimate trait-specific heritability enrichment while accounting for LD (Finucane et al. 2015; Speed and Balding 2019), these approaches are optimized for analyses involving large sets of variants and tend to lose power when applied to smaller SNP subsets. Given the limited size of our ASAS-associated SNP set, we adopted an alternative strategy to evaluate their potential relevance to disease. Specifically, we grouped GWAS traits into biologically meaningful broad categories, including AD-related, brain-related, and immune-related traits, and assessed whether ASAS-associated SNPs were enriched for LD with GWAS loci from each category (Methods). This targeted approach revealed clear enrichment for ASAS-associated SNPs among loci linked to AD- and brain-related traits, while showing minimal overlap with general traits such as height and BMI (Figure 3C). Notably, we also observed enrichment in immune-related traits, aligning with prior evidence that immune signaling contributes to both brain function and AD pathology (Castellani et al. 2023; Jorfi et al. 2023).

Within the AD-related GWAS terms, we observed that over 50% of ASAS-associated SNPs in LD with GWAS loci resided in AD-relevant genes (Rath et al. 2021) (Figure 3D), suggesting that their splicing effects may underlie a portion of the observed genetic risk. Additionally, six of our ASAS-associated SNPs were themselves direct AD GWAS hits, accounting for ∼14% of all direct GWAS overlaps observed in our dataset. Together, these findings reinforce the important role of splicing regulation in AD biology.

### AD-specific ASAS patterns

Given the observed enrichment of ASAS events in AD-relevant loci and genes, we next asked whether specific tag SNPs exhibited differential allelic bias between AD and control samples. To address this, we adapted a statistical framework previously developed in our lab for group comparisons of proportion data in sequencing-based analyses (Tran et al. 2020) (Supplemental Fig. S4A; Methods). Briefly, this method models allelic counts using a beta-binomial distribution, which captures both biological variability across samples and technical uncertainty due to sequencing depth. It then tests for differences in allelic bias between two groups by comparing a null model assuming a shared distribution across all samples with an alternative model allowing group-specific distributions. To distinguish robust biological shifts from experimental noise (Serre et al. 2008), we imposed a minimum mean allelic ratio difference of 0.1 between AD and controls. Given this stringent effect size criterion, we applied a nominal p-value cutoff of 0.05 rather than an FDR threshold to identify differential events.

Across all brain regions, we identified 54 unique tag SNPs within 50 genes that exhibited significant differences in allelic ratios between AD and control samples (Supplemental Table S3), with the number of events varying by region (Supplemental Fig. S4B, C). Several of these SNPs were located in genes previously implicated in AD (Figure 4A), indicating potential disease-specific regulation of splicing. Notably, SNPs in *CNTN1* and *SLC6A1* showed some of the most pronounced shifts in mean allelic ratios, highlighting two distinct pathological mechanisms. *CNTN1*, a regulator of synapse formation and neuroinflammatory (Li et al. 2021), is known to interact with the amyloid-β precursor protein (APP) (Bai et al. 2008). Its expression is upregulated in postmortem AD brains, and its overexpression in mouse models drives cognitive deficits (Li et al. 2023). Conversely, the GABA transporter gene *SLC6A1* is downregulated in postmortem AD brains, reflecting significant disruption to inhibitory signaling pathways (Fuhrer et al. 2017; Sun et al. 2023; Carello-Collar et al. 2023).

**Figure 4.**
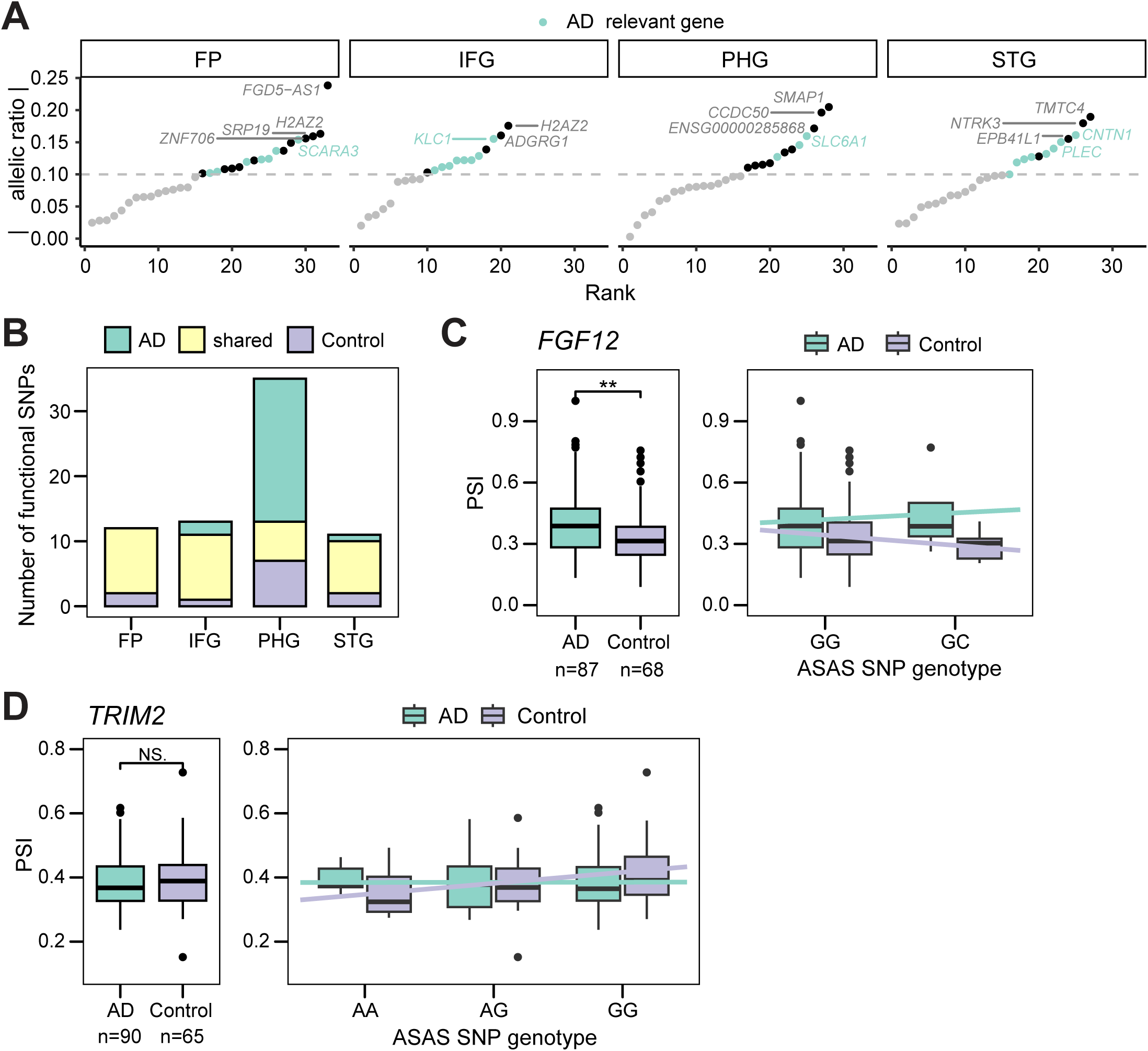
AD-specific ASAS patterns. (A) SNPs exhibiting differential allelic ratios between AD and control groups in different brain regions, ranked by their absolute difference in mean allelic ratio (Methods). Gene names are shown if the associated SNPs had an absolute difference greater than 0.15. Significant SNPs located in AD-relevant genes are colored green. STG: superior temporal gyrus; PHG: parahippocampal gyrus; IFG: inferior frontal gyrus; FP: frontal pole. Ensembl IDs are shown for genes without approved HGNC symbols. (B) Overlap of putative functional SNPs that were testable in both AD and control groups in each brain region (Methods). (C) and (D) Percent-spliced-in (PSI) values for an ASAS exon associated with a functional SNP within *FGF12* (C) and *TRIM2* (D), separated into AD and control groups (left) or by the genotype at the functional SNP (right). Statistical significance of the PSI between AD and control groups was evaluated with a Wilcoxon Rank-sum test. ***p < 0.001. N.S.: not significant.

The differential allelic ratio analysis described above relies on RNA-seq coverage, which inherently excludes functional SNPs located in intronic regions. To encompass the full set of candidates (both intronic and exonic), we adopted a different strategy: identifying SNPs that were genetically testable in both AD and control groups but were predicted as ‘functional’ in only one. (Figure 4B). For example, in the *FGF12* gene, we identified a functional SNP (rs62293022) in the parahippocampal gyrus exhibiting opposing allele-specific trends in exon inclusion between AD and control samples (Figure 4C). This SNP was associated with overall increased exon inclusion in AD cases. Notably, prior studies reported significant downregulation of *FGF12* in the parahippocampal gyrus of patients with AD (Britton et al. 2023). This gene encodes a fibroblast growth factor that interacts with voltage-gated sodium channels (Wildburger et al. 2015; Goldfarb et al. 2007) and is linked to epilepsy (Tian et al. 2021), a condition known to co-occur with AD (Kamondi et al. 2024).

A similar AD-specific splicing pattern was observed for *TRIM2*, a gene essential for neuronal growth and regulation (Lokapally et al. 2020). Specifically, heterozygous control samples at the SNP rs4696445 showed increased exon inclusion relative to homozygous samples, a trend that was notably absent in the AD samples (Figure 4D). This loss of regulation is significant given that *TRIM2* deficiency in mice leads to neurofilament light subunit accumulation, axonopathy, and progressive neurodegeneration (Balastik et al. 2008). Additionally, *TRIM2* plays an antiviral role in both murine models (Sarute et al. 2019) and human dermal fibroblasts (Kimsa et al. 2014), which is particularly relevant given the hypothesis that viral activation may contribute to AD etiology (Itzhaki et al. 2016). Neither of the identified SNPs were detected as sQTLs in GTEx brain tissues, emphasizing the added value of our approach in uncovering context-specific genetic regulation of splicing that may be missed by conventional methods. Collectively, these results highlight the functional significance of ASAS-associated SNPs in AD and support their role in modulating splicing in a disease-specific manner.

### 5’ UTR splicing events affect protein abundance and isoforms

We analyzed the genomic distribution of unique ASAS events and observed a significant enrichment in the 5’ UTR of transcripts (Figure 5A), a pattern consistently observed across all four brain regions (Supplemental Fig. S5). Given the critical role of the 5’ UTR in regulating mRNA stability (Jia et al. 2020) and translation efficiency (Calvo et al. 2009), we hypothesized that ASAS events in these regions influence the proteome through three primary mechanisms: (1) modulating RNA expression (leading to changes in overall protein abundance), (2) remodeling protein isoforms (e.g., via alternative start codons), or (3) tuning protein abundance without protein isoform changes (e.g., via regulating translational efficiency).

**Figure 5.**
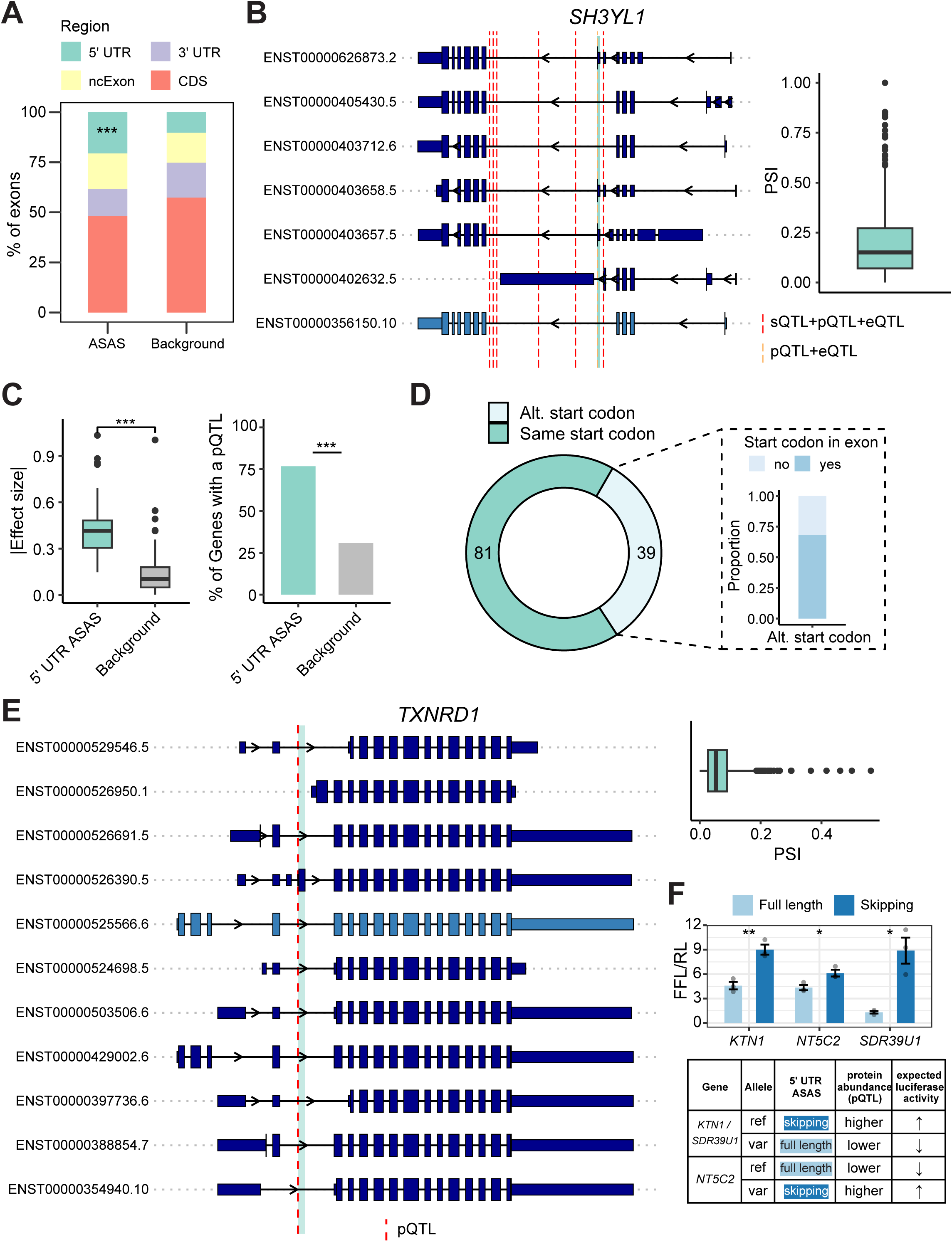
5’ UTR splicing events may affect protein abundance and isoforms. (A) Genomic distribution of unique ASAS exons compared to all annotated exons as a background. ***p = 1.28 x 10^-13^ (Fisher’s exact test). CDS: coding sequence. ncExon: exons in non-coding transcripts. (B) Transcript isoforms for *SH3YL1* from the Gencode basic annotation, with the 5’ UTR ASAS exon highlighted in green (left). The light blue transcript indicates the Ensembl canonical transcript. The red dashed line denotes the location of a putative functional SNP overlapping a pQTL. Taller rectangles represent protein-coding exons and shorter rectangles represent UTR exons. Right: PSI values of the 5’ UTR ASAS exon across samples. The orange and red dashed lines denote the locations of ASAS-associated SNPs overlapping with a pQTL and an eQTL or a pQTL, an sQTL, and an eQTL, respectively. (C) Left: Absolute effect size of 5’ UTR ASAS-associated SNPs overlapping pQTLs. The same number of random pQTLs were sampled as the background distribution. The significance was evaluated using a Wilcoxon Rank-sum test. ***p < 0.001. Right: Percentage of genes harboring 5’ UTR ASAS events that contain at least one pQTL. The background percentage was calculated using all annotated genes. ***p < 2.2 x 10^-16^ (Fisher’s exact test). (D) Number of genes with transcripts containing 5’ UTR ASAS exons utilizing the same or an alternative start codon relative to the isoforms where the ASAS exons are skipped (top), and the proportion of the alternative start codons that fall within the ASAS exon itself (right). (E) Similar as (B), but for the gene *TXNRD1*. The red dashed line denotes the location of a putative functional SNP overlapping a pQTL. (F) Top: Relative luciferase activity (Firefly/Renilla Luciferase, FFL/RL) comparing protein abundance between full-length and 5’ UTR exon-skipping isoforms for *KTN1*, *NT5C2*, and *SDR39U1*. Bottom: Concordance between allele-specific splicing and pQTL directionality. The table maps each allele (reference vs. variant) to its associated isoform and observed protein abundance trend (pQTL) to define the expected direction of luciferase activity. Significance was determined by a two-sided t-test (* P < 0.05, ** P < 0.01).

To establish a baseline for these mechanisms, we first evaluated the concordance between mRNA abundance and protein levels using matched proteomics data (Bai et al. 2020) (Methods). Genome-wide, only ∼33% of genes showed a significant correlation between mRNA and protein abundance (Supplemental Fig. S6A). However, this concordance was significantly higher (∼47%, p = 0.01, Fisher’s Exact test) for genes containing 5’ UTR ASAS events. This suggests that for a substantial subset of these candidates, the splicing variation may be intrinsically coupled to mRNA abundance, thereby driving a concordant change in protein levels.

To test this “abundance-associated” mechanism, we examined the overlap between 5’ UTR ASAS-associated SNPs and expression quantitative trait loci (eQTLs) from GTEx brain tissues and pQTLs from a recent multi-ancestry large-scale AD and brain proteomic study (Wingo et al. 2025). We identified 20 SNPs that overlapped both pQTLs and eQTLs, supporting a model where the splicing event co-occurs with shifts in mRNA availability (Supplemental Fig. S6B). A clear example is *SH3YL1*, which encodes a protein containing an SH3 domain and has been implicated in dorsal ruffle formation and interactions with cytoskeletal-related proteins (Hasegawa et al. 2011), but little is known about its potential brain-related functions. Within this gene, seven ASAS-associated SNPs overlapped both pQTLs and eQTLs, and six of these were also identified as sQTLs in GTEx brain tissues (The GTEx Consortium 2020) (Figure 5B), highlighting their potential role in regulating mRNA abundance, splicing, and protein abundance. The associated 5’ UTR ASAS exon exhibited highly variable inclusion levels across individuals, with PSI values ranging from 0% to nearly 100%, suggesting strong inter-individual differences in splicing regulation. *SH3YL1* has been previously linked to neurological traits; for instance, a GWAS study of suicide severity in bipolar disorder identified a risk locus encompassing this gene (Zai et al. 2015), and its potential roles in brain function and AD have been proposed (Seyfried et al. 2011). More recently, Bayesian fine-mapping analyses have identified *SH3YL1* as a strong causal mediator of AD risk (Park et al. 2017), further underscoring the functional importance of this splicing event in the brain.

However, the majority of our findings point to a second, “isoform-remodeling” mechanism where protein isoforms are altered without necessarily changing total mRNA abundance. We identified 103 ASAS-associated SNPs that overlapped with pQTLs and found that these SNPs had greater absolute effect sizes compared to randomly sampled pQTLs (Figure 5C, left). Furthermore, genes harboring 5’ UTR ASAS events were significantly enriched for pQTLs, with 76% of these genes containing at least one pQTL, in contrast to only ∼30% of genes in the background set (Figure 5C, right). Strikingly, >80% of the 5’ UTR ASAS-associated SNPs that overlapped a pQTL did *not* overlap an eQTL (Supplemental Fig. S6B), implying that their direct impact on protein isoform expression.

To further explore whether 5’ UTR ASAS events were associated with altered protein isoforms, we examined transcript annotations to compare the start codon positions between isoforms that include versus skip the ASAS exon. Notably, 39 of the 120 transcripts (approximately 30%) containing the 5’ UTR ASAS exon utilized an alternative start codon relative to the skipping isoform, with the majority of these alternative start codons located within the ASAS exon itself (Figure 5D). Among these, 15 ASAS exons were directly linked to SNPs overlapping pQTLs, including two within both mitochondrial (Rath et al. 2021) and AD-related genes *TXNRD1* and *MFF*, suggesting another potential mechanism by which ASAS events in the 5’ UTR influence both protein abundance and protein isoform composition.

For instance, in the *TXNRD1* gene, one transcript isoform (ENST00000526390.5) contains a 5’ UTR ASAS exon that introduces an alternative start codon located within the exon itself, while isoforms that skip the exon initiate translation from different positions (Figure 5E). Although this exon generally exhibits low inclusion across samples, a subset of 81 showed markedly higher inclusion levels (PSI > 15%). A putative functional SNP located at the second base of the ASAS exon directly overlaps with a pQTL and not an eQTL in brain tissue, suggesting that this variant may influence both exon inclusion and protein abundance. Notably, inclusion of this ASAS exon leads to an isoform that lacks a portion of the glutaredoxin domain found in the canonical isoform (Paysan-Lafosse et al. 2025), potentially altering its redox-related activity. *TXNRD1* is involved in the mitochondrial proteome (Rath et al. 2021) and plays a critical role in mitigating oxidative stress (Arnér 2009), a process closely linked to inflammation, aging (Finkel and Holbrook 2000), and neurodegeneration (Butterfield and Halliwell 2019). Moreover, genetic variants in *TXNRD1* have been associated with traits such as longevity and cognitive performance (Dato et al. 2015, 2014; Soerensen et al. 2012), further supporting its relevance to brain aging and disease.

Additionally, the 81 ASAS events that do not affect the primary start codon may influence protein abundance through a distinct mechanism, such as modulating translational efficiency. To test this hypothesis, we cloned the full-length and exon-skipping 5’ UTR isoforms of 3 candidate genes (*KTN1, NT5C2*, and *SDR39U1*) upstream of a luciferase reporter (Methods). In all three cases, we observed significant isoform-dependent differences in reporter activity (Figure 5F, top). Importantly, these experimental results are consistent with our prediction: the specific allele associated with the isoform (skipping or inclusion) that drove higher luciferase activity corresponded to the allele associated with increased protein abundance in pQTL studies (Figure 5F, bottom). Interestingly, the functional SNP with allele-specific binding to QKI (Figure 2E) was predicted to regulate the same 5’ UTR *KTN1* exon here, further highlighting the interplay between the multiple regulatory layers. These findings experimentally validate that 5’ UTR ASAS serves as a mechanism for pQTL effects, regulating protein abundance independent of coding sequence alterations.

## Discussion

In this study, we present a systematic map of ASAS in the human brain, nominating hundreds of functional regulatory variants that would be invisible to standard differential expression analyses. Beyond cataloging these events, our work reveals two fundamentally new insights into how splicing variation shapes the AD proteome. First, we identify 5’ UTR splicing as a critical determinant of the proteome, acting through dual mechanisms: while a subset of events modulates total protein abundance by altering mRNA levels (e.g., *SH3YL1*), the majority drive isoform remodeling without changing transcript abundance. Second, we uncover a class of “disease-context-specific” variants that drive opposing splicing patterns in AD patients compared to controls, rewiring the regulatory logic of key genes like *TRIM2* and *FGF12* in the presence of pathology. By linking these events to specific RBP motif disruptions (e.g., QKI, TIA1), we establish ASAS not just as a source of transcript diversity, but as a functional regulatory layer that directly shapes the brain proteome and disease risk.

A major finding of our work is the functional dominance of splicing events in the 5’ UTR. While previous studies have linked splicing to protein traits (Abood et al. 2024; Tokolyi et al. 2025; Pietzner et al. 2021), the contribution of 5’ UTR splicing remains poorly characterized. We found that these events frequently overlap with brain pQTLs and may mechanistically drive protein variance by modulating mRNA levels (those that also overlap with eQTLs), translation efficiency or alternative translation start sites. Most transcriptomic and proteomic studies focus on coding regions, overlooking how variation in the 5’ UTR may affect translation initiation, isoform diversity, and gene regulation. Our results address this gap by highlighting (supported by experimental validation) allele-specific splicing in 5’ UTRs as a driver of protein abundance independent of coding changes, particularly in the brain. These findings suggest that 5’ UTR splicing should be more systematically studied, as it may represent an underappreciated link between noncoding genetic variation, post-transcriptional regulation, and disease phenotypes such as AD.

Biologically, these ASAS events converged on pathways central to neurodegeneration. We observed a strong enrichment in genes involved in mitochondrial function, particularly respiratory chain complex I assembly, a pathway tightly linked to aging, neurodegeneration, and specifically AD pathology (Rhein et al. 2009; Aksenov et al. 1999). While mitochondrial dysfunction is a known hallmark of AD (Fairbrother-Browne et al. 2021; D’alessandro et al. 2025; Holper et al. 2019), our findings add a new dimension by implicating allele-specific splicing as a regulatory layer influencing inter-individual differences in mitochondrial activity.

In addition to broad patterns of splicing regulation, we identified ASAS events exhibiting disease-specific differences between AD and control samples. These included shifts in allelic bias at known AD genes such as *CNTN1* and *SLC6A1*. Furthermore, by comparing functional SNP predictions across disease groups, we uncovered ASAS events specific to AD brains in genes like *FGF12* and *TRIM2*, both of which are linked to AD risk and neurological phenotypes. Notably, these ASAS events were not detected in sQTL datasets, underscoring the added resolution of allele-specific analyses in uncovering disease-context splicing regulation. These results emphasize the value of incorporating disease stratification when analyzing splicing patterns and highlight ASAS as a potential mechanism through which genetic variation contributes to AD-specific molecular alterations. Future experimental validation will be important to confirm these effects, although it will require specialized AD-relevant cellular context and models to capture disease-specific alterations that simple reporters do not often recapitulate.

To facilitate the interpretation of these biological findings, we used a concordance-based approach (Amoah et al. 2021) to nominate over 500 putative functional SNPs. These variants were supported by multiple lines of evidence, including close proximity to regulated exons, overlap with brain sQTLs, and enrichment for RBP binding sites involved in splicing regulation. Importantly, they were significantly enriched in GWAS loci linked to AD, brain function, and immune traits, offering immediate functional hypotheses for associations observed in complex trait studies. By pinpointing variants that physically disrupt splicing motifs, this candidate set functions as a “fine-mapping” resource for the community, prioritizing SNPs most likely to have a causal regulatory impact in the brain.

Our study has limitations inherent to the study design and available data. First, our concordance-based approach is constrained by sample size, read length, sequencing depth, and genotype availability. The usage of short-read (100bp) RNA-seq limits our ability to detect complex multi-exon events, intron retention, or events in low-complexity regions, though standard cassette exons are well captured. Second, cell-type specific splicing may not be captured in bulk RNA-seq. For example, the well-characterized AD splicing variant in *CD33* (Hollingworth et al. 2011; Naj et al. 2011; Malik et al. 2013) was not detected due to low expression the gene in our bulk tissue samples (RPKM < 0.3). Third, our mechanistic predictions rely on eCLIP data from HepG2 and K562 cell lines, which may overlook brain-specific RBP interactions. The necessity of studying the relevant tissue is highlighted by the minimal overlap we observed between our brain ASAS variants and regulatory variants identified in blood cells (Astle et al. 2016), reinforcing the distinct nature of the brain regulatory landscape. Finally, our method relies on allelic ratios in mature mRNA, thus cannot capture splicing events that trigger nonsense-mediated decay (NMD) and biases all allelic ratios across the gene. Despite these limitations, our results highlight the utility of splicing-resolved analyses for interpreting genetic variation.

In summary, our study provides a comprehensive map of allele-specific alternative splicing in the human brain, identifying regulatory variants that drive transcript diversity, protein isoform remodeling, and disease risk. By integrating transcriptomics and proteomics, we demonstrate that 5’ UTR splicing is a key mechanism shaping AD proteome. Future work extending these analyses to single-cell and long-read sequencing contexts will be essential to fully elucidate how these splicing alterations contribute to the cell-type-specific dysfunction observed in neurodegenerative disease.

## Methods

### Identification of allele-specific alternative splicing events

We downloaded RNA-seq alignment files in bam format and WGS-based variants in VCF format from the Mount Sinai Brain Bank (MSBB) cohort (Wang et al. 2018). Bam files were converted to FASTQ format with samtools bam2fq and realigned to the hg38 genome using STAR(Dobin et al. 2013) with the ENCODE standard option parameters. We discarded samples based on the exclusion criteria (Wang et al. 2018) described by MSBB and excluded samples without genotype information, resulting in 1,047 samples across 4 brain regions from 293 donors.

To harmonize the genome coordinates of genotype data, VCF files were first converted from GRCh37 (b37) to hg19, then from hg19 to hg38 using GATK’s LiftoverVCF tool. We retained only heterozygous SNPs labeled “PASS” in the VCF file for downstream analysis. Using these high-confidence heterozygous variants, we identified alternative mRNA processing events associated with allele-specific expression of nearby SNPs in each sample, following our previously published method^17^. Briefly, this approach compares the allelic ratios of SNPs located within alternatively spliced exons to those in constitutively spliced exons of the same gene, enabling the identification of ASAS events in each sample. SNPs showing a significant allelic imbalance specific to the alternative region were designated as *tag SNPs*, which serve as markers for downstream functional SNP prediction (see below). We removed ASAS events predicted in HLA genes from further analyses due to the highly polymorphic nature of these genes.

### Gene ontology (GO) enrichment analysis

We performed GO (Ashburner et al. 2000; The Gene Ontology Consortium et al. 2023) enrichment using annotations from the Ensembl (Harrison et al. 2024) database and the R package biomaRt (Durinck et al. 2009). For each query gene, we defined a matched background gene by selecting genes with similar characteristics: expression level within ±10%, gene length within ±10%, and exclusion of genes containing ASAS events. To assess enrichment significance, we selected 10,000 sets of random background genes for the query gene set and calculated the frequency of each GO term in these sets to generate a null distribution. The empirical p-value for each GO term was calculated based on its frequency in the query set relative to the background distribution, and significance was determined using a false discovery rate (FDR) threshold of 0.05.

### Prediction of functional SNPs using concordance scoring

To identify putative functional SNPs that regulate ASAS, we implemented a concordance-based approach as described in our previous work (Amoah et al. 2021). This method evaluates how well the genotype of a candidate SNP explains the allelic imbalance observed at a tag SNP associated with an ASAS event. Briefly, we first defined the allelic imbalance at a tag SNP as *d*_*i*_ = |0.5 − *R*_*i*_|, where *R*_*i*_ is the allelic ratio of the 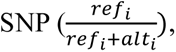 and *ref*_*i*_ and *alt*_*i*_ are the read counts for the reference and alternative allele, respectively. The concordance score between a candidate SNP and a tag SNP was then defined as below:

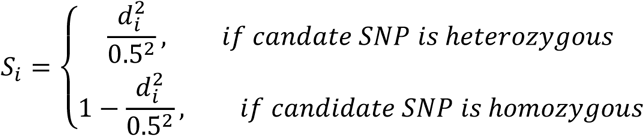

If the candidate SNP being tested is the same as the tag SNP, then 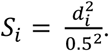

We applied this framework to all common dbSNPs in the same gene as a given tag SNP, generating a distribution of *S*_*i*_ scores across the population for each candidate SNP. For a truly functional SNP, the distribution of *S*_*i*_ scores is expected to exhibit either (1) a single peak close to 1, when the SNP causes a strong splicing difference between alleles, or (2) two peaks - one near 1 for homozygous individuals and another intermediate peak contributed by heterozygous individuals. In contrast, non-functional SNPs are expected to have a distribution skewed toward 0.

To identify candidate functional SNPs, we tested whether the *S*_*i*_ score distribution deviated significantly from the null expectation and whether it was supported by sufficient sample representation. Specifically, we required that: at least 10% of individuals were heterozygous for the candidate SNP (based on previous simulation benchmarks) (Amoah et al. 2021). Additionally, the total number of *S*_*i*_ values in the candidate-tag SNP pair exceeded 10% of the total sample size within a given region and disease condition.

Prefrontal cortex samples were excluded from this analysis due to low sample size (n = 13). This concordance-based scoring method enables prioritization of likely functional *cis*-regulatory variants that underlie observed allele-specific splicing patterns.

### Estimation of brain region and individual contributions to ASAS variation

To quantify the relative contributions of individual and region-specific factors to ASAS variation, we modeled allelic imbalance for each ASAS exon using a linear mixed-effects model implemented via lmer function from the lme4 R package (version 4.2.2 (R Core Team (2022))). The model was specified as follows:

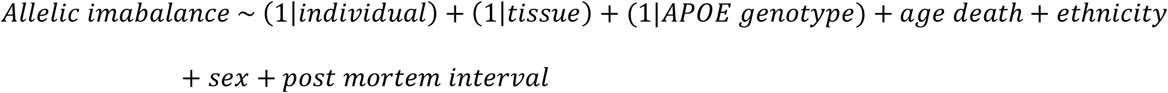

Random effects for individual, brain region (tissue), and *APOE* genotype were included to estimate the variance explained by these factors, while fixed effects (age of death, sex, ethnicity, and postmortem interval) were chosen based on previous literature (Amoah et al. 2021; Melé et al. 2015). Variance components derived from the model were used to assess the relative impact of individual, brain region, and *APOE* genotype contributions on ASAS patterns.

### RBP binding site enrichment and binding affinity analysis

To investigate whether functional SNPs affect RBP regulation, we assessed their enrichment in experimentally defined RBP binding sites. eCLIP data for RBPs in HepG2 and K562 cell lines were obtained from the ENCODE Consortium (Van Nostrand et al. 2020a). We used GREGOR (Schmidt et al. 2015) (Genomic Regulatory Elements and Gwas Overlap algoRithm) to test whether functional SNPs and their LD proxies overlapped RBP binding sites more frequently than expected by chance. GREGOR compares the observed overlap to an empirical null distribution generated from ∼500 matched control SNPs, using EUR LD reference data from the 1000 Genomes Phase 1 v2 Panel (Durbin et al. 2010) with an LD threshold of *R^2^* ≥ 0.8 and a 200 kb window. RBPs with enrichment at a Benjamini-Hochberg FDR ≤ 0.05 were considered significant.

For significantly enriched RBPs, we next evaluated whether functional SNPs altered RBP binding affinity. Using pre-trained models from DeepRiPe (Ghanbari and Ohler 2020), we computed the absolute change in predicted binding scores between the reference and alternative alleles of each functional SNP. To determine if these changes were greater than expected, we sampled the same number of common SNPs from the dbSNP database (excluding genes with functional SNPs) and repeated this sampling 1,000 times. For each iteration, we computed the median change in binding affinity score for each RBP and compared the distribution across 1,000 iterations to the observed median change from the functional SNPs. Fold change was defined as the ratio of the median change for functional SNPs over the median from random SNPs. Statistical significance of the fold change was evaluated with a one-sided Mann-Whitney U test, with a null hypothesis that the fold change equals 1.

### Overlap of GTEx sQTLs with functional SNPs

We downloaded significant sQTLs in brain tissues from the GTEx V8 dataset (https://www.gtexportal.org/home/downloads/adult-gtex/qtl) and intersected their genomic coordinates with those of the functional SNPs we identified. To assess enrichment, we additionally downloaded all tested variant-gene cis-sQTL associations from GTEx V8 across brain tissues. For each brain region, we identified functional SNPs that overlapped sQTLs with shared exon coordinates. We then compared the distribution of nominal p-values from these overlapping sQTLs to those from a matched background set, generated by randomly sampling the same number of SNPs from all tested cis-sQTL. Enrichment was calculated using the Kolmogorov-Smirnov test.

### Enrichment of ASAS-associated SNPs in GWAS loci

We assessed whether ASAS-associated SNPs were enriched in GWAS loci using data from the GWAS catalog (Buniello et al. 2019) (version 1.0.2, downloaded 04-April-2024). The dataset was filtered to include only associations with p-values less than 5.0e-8. GWAS SNPs were grouped into LD blocks based on CEU population data, with an *R^2^* ≥ 0.8 and *D’* ≥ 0.9. For each ASAS-associated SNP, we determined whether it was in LD with at least one GWAS SNP and within a 200 kb window. To establish a background, we randomly sampled the same number of common SNPs from the dbSNP database, restricted to genes without ASAS events. A Fisher’s exact test was used to compare the number of ASAS-associated SNPs versus background SNPs directly overlapping a GWAS SNP.

To assess trait-specific enrichment, we calculated an enrichment score of each GWAS term by comparing the proportion of ASAS-associated SNPs in LD with the GWAS SNPs for that trait to the proportion from randomly sampled common SNPs. The enrichment score was defined as:

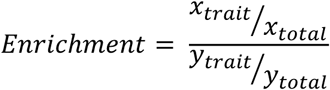

where:

*x*_*trait*_ = number of ASAS-associated SNPs in LD with GWAS SNPs for the trait

*x*_*total*_ = total number of ASAS-associated SNPs in LD with at least one GWAS SNP

*y*_*trait*_ = number of random dbSNPs in LD with GWAS SNPs for the trait

*y*_*total*_ = total number of random dbSNPs in LD with at least one GWAS SNP

To enable interpretation at the trait-category level, we grouped the GWAS terms containing “Alzheimer”, “dementia”, “amyloid”, and “neurofibrillary tangles” into an “AD-related” category. Similarly, terms including “neuroticism”, “Parkinson”, “schizophrenia”, and “brain” were grouped into an “Other brain-related” category. “Immune-related” traits were defined by terms containing “immune”, “antigen”, “microglia”, “cytokine”, “antibody”, “interferon”, or “inflammatory”. Fisher’s exact tests were used to evaluate the statistical significance of enrichment for each category.

### Identification of allelic imbalance differences between AD and control samples

To identify SNPs exhibiting differential allelic ratios between AD and control samples, we applied a beta-binomial modeling framework previously developed for detecting differential RNA editing, which is generally applicable to model the allelic ratio of SNPs and infer the difference between two groups (Tran et al. 2020) (Supplemental Fig. S4A). This model accounts for both biological variability and read coverage when estimating differences in allelic ratios between groups. A SNP was considered testable if it was previously identified as a tag SNP for ASAS in at least 5 AD and 5 control samples for a given brain region. We identified significant differential allelic imbalance using a p-value cutoff of 0.05 and required the minimum absolute difference in mean allelic ratios to be at least 0.1 between AD and control groups, as differences below this threshold may be due to experimental variability (Serre et al. 2008).

### mRNA and protein abundance correlation

We downloaded the MSBB TMT proteomics mean-centered, corrected matrix (Bai et al. 2020) using Synapse (syn24983526) and calculated the Spearman correlation between RPKM and protein abundance values for each gene. We calculated the correlation for 186 parahippocampal gyrus samples that had both protein abundance and RPKM information. Read counts per gene were obtained using the HTSeq software (Putri et al. 2022) and RPKM values from the edgeR package (Chen et al. 2025). Samples with an RPKM of 0 were removed from correlation calculations for a given gene. Significant correlations were determined at an FDR cutoff of 0.05.

### Overlap with pQTLs

We assessed the overlap between ASAS-associated SNPs and pQTLs using data from a multi-ancestry large-scale proteomic study (Wingo et al. 2025). This study included samples collected from multiple sites, including Mount Sinai University Hospital. We used significant pQTLs (FDR ≤ 0.05) from all ancestries to evaluate whether 5’ UTR ASAS-associated SNPs overlapped pQTLs at the SNP level and whether their corresponding genes contained at least one significant pQTL. To assess statistical significance, we generated a control distribution by randomly sampling the same number of pQTLs from the full dataset and repeated the sampling procedure to establish an empirical background for SNP-level overlap.

### Purification of recombinant human ΔQKI

PCR-amplified cDNA fragments encoding the STAR domain of human QKI (residues 7–214) were cloned into the pET-28a vector downstream of an N-terminal 6×His tag followed by a thrombin cleavage site. The primers used are listed in Supplemental Table S4. The construct was transformed into Rosetta (DE3) competent cells (Novagen). Protein expression was induced with 0.5 mM isopropyl β-D-1-thiogalactopyranoside (IPTG) at an OD₆₀₀ of ∼1.0, followed by incubation at 18 °C for 18 h with shaking at 180 rpm. Cells were harvested by centrifugation at 7,000 × g for 5 min at 4 °C and resuspended in 10 mL ice-cold lysis buffer A (20 mM HEPES, pH 8.0, 150 mM NaCl, 20 mM imidazole, 1 mM TCEP, PMSF). Cells were lysed by sonication at 30% amplitude with 10-s on/3-s off pulses for 30 min. The lysate was clarified by centrifugation at 7,000 × g for 30 min at 4 °C, and the supernatant was filtered through a 0.22 μm syringe filter. The filtered lysate was loaded onto Ni-NTA resin (Thermo Fisher Scientific), washed with buffer A containing 50 mM imidazole, and eluted with buffer A supplemented with 500 mM imidazole. The eluted protein was further purified by size-exclusion chromatography using a Superdex 200 10/300 GL column. Protein purity was assessed by SDS-PAGE, and protein concentration was determined using BioDrop spectrophotometer.

### In vitro transcription of ΔQKI target RNA

DNA templates (45 bp) for ΔQKI target RNAs were synthesized as oligonucleotides (IDT). Double-stranded transcription templates were generated by annealing forward and reverse oligonucleotides at a 1:1 molar ratio. Sequences of the DNA templates and the corresponding RNAs are listed in Supplemental Table S4. In vitro transcription was performed at 37 °C for 4 h using the HiScribe T7 Quick High Yield RNA Synthesis Kit (NEB). Transcribed RNAs were purified by 15% denaturing polyacrylamide gel electrophoresis (7 M urea), followed by extraction using the RNA Clean & Concentrator-5 Kit. RNAs were subsequently biotinylated at the 3′ end using the Pierce RNA 3′ End Biotinylation Kit to attach a single biotinylated nucleotide. Biotinylated RNAs were purified again by urea-PAGE and RNA Clean & Concentrator-5 Kit. RNA concentrations were measured using BioDrop spectrophotometer.

### Electrophoretic Mobility Shift Assay (EMSA)

Purified biotinylated RNA probes (30 pmol) were incubated with recombinant ΔQKI protein (0, 0.4, 1.0, or 2.0 μg) in a total volume of 20 μL binding buffer (20 mM HEPES, pH 8.0, 150 mM NaCl, 1 mM TCEP, 0.1× protease inhibitor cocktail, and 10 U RNase inhibitor) at room temperature for 30 min. Samples were resolved on 5% TBE-PAGE gels at 75 V for 1.2 h. RNAs were transferred to positively charged nylon membranes using a Trans-Blot Turbo Transfer System for 30 min, followed by UV cross-linking at 1,250 mJ/cm². Membranes were blocked in 3% BSA prepared in 1× TBS with 0.1% Tween-20 for 30 min at room temperature, then incubated with streptavidin-HRP (Thermo Fisher Scientific) for 30 min. Membranes were washed three times with TBST for 5 min each and developed using ECL reagents (Cytiva). Signals were visualized using an iBright 750 imaging system. Band intensities were quantified using ImageJ, with the intensity from the 2.0 μg ΔQKI condition normalized to 1.

### Dual luciferase reporter assay

To investigate the translational effects of ASAS exons within 5’ UTRs, genomic DNA from K562 cells was used as a template to PCR-amplify the corresponding full-length 5’ UTR sequence of *NT5C2*, *KTN1* or *SDR39U1*. The resulting fragments were cloned into the luciferase reporter vector pGL3-basic (Promega) using Gibson Assembly (NEBuilder HiFi DNA Assembly, NEB). Exon-skipping constructs were generated by whole-plasmid PCR-based mutagenesis. The pRL-TK plasmid was used as an internal reference. The full length and skipping sequences are listed in Supplemental Table S4. HEK293T cells were seeded at a density of 0.5 million cells per well in 12-well plates and allowed to attach overnight. Cells were co-transfected with 2 μg of pGL3-Firefly plasmid and 0.3 μg of pRL-Renilla control plasmid using Lipofectamine 3000 (ThermoFisher). After 24 hours of incubation, cells were lysed in 200 μl of 1X passive Lysis buffer (25 mM Tris phosphate pH 7.8, 4 mM EGTA, 1% Triton X-100, 10% glycerol, 2 mM dithiothreitol, 1.25 mg/ml lysozyme, and 2.5 mg/ml BSA). Firefly luciferase activity was measured using a VICTOR Nivo plate reader (PerkinElmer) following the addition of 100 μl of Luciferase Assay Substrate (Promega) to 20 μl of cell lysate. Subsequently, 100 μl of Stop & Glo reagent (Promega) was added to quantify Renilla luciferase activity. Relative luciferase activity was calculated by normalizing Firefly luminescence (FFL) to Renilla luminescence (RL). All experiments were performed in three independent biological replicates.

## Acknowledgements

We thank members of the Xiao laboratory for helpful discussions and comments on this work. This work was supported in part by grants from the National Institute of Health (R01AG075206, and R01AG056476).

## Author Contributions

A.J.K., K.A., and X.X. designed the study. A.J.K., K.A., R.M.B, and G.Q.V. performed bioinformatic analyses. L.Z. and J.H.B. performed the EMSA and luciferase experiments. A.J.K. and X.X. wrote the original draft. All authors reviewed and approved the final manuscript.

