## Supplemental Figures for "Allele-specific splicing modulates protein isoforms and Alzheimer’s risk"

### Figure S1

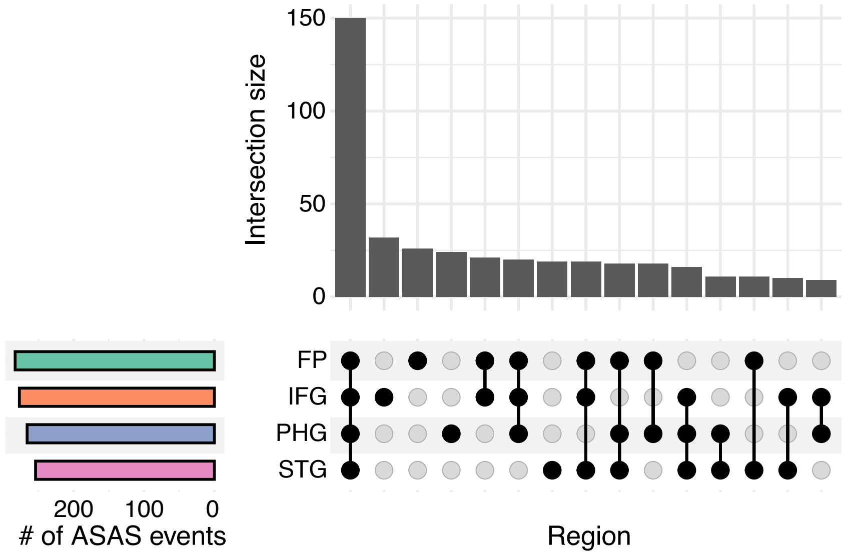


**Figure S1. Overlap of testable ASAS events across brain regions.**
Overlap of ASAS events that were testable in at least one sample per brain region across brain regions. STG: superior temporal gyrus; PHG: parahippocampal gyrus; IFG: inferior frontal gyrus; FP: frontal pole.

### Figure S2

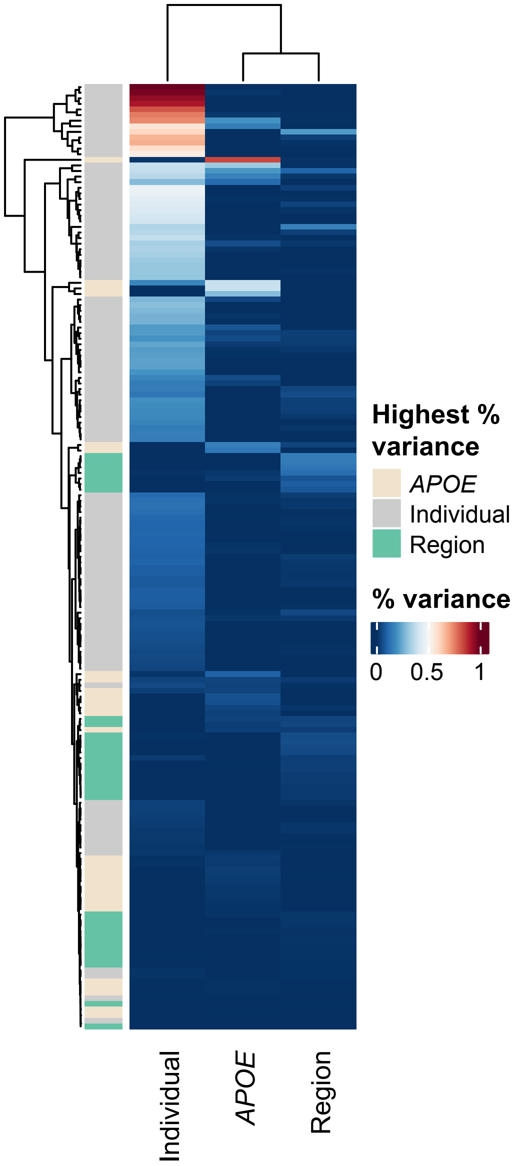


| % variance from APOE | Gene | Chromosome | AS_region |
|---|---|---|---|
| 84.04% | *VSTM2A-OT1* | chr7 | 54557850:54559971 |
| 38.65% | *FAM86FP* | chr12 | 8234276:8234426 |
| 38.47% | *ANKRD36* | chr2 | 97163244:97163854 |
| 28.36% | ***RNF19A*** | chr8 | 100300528:100300651 |
| 15.73% | *SNHG14* | chr15 | 25204684:25204728 |
| 13.90% | *ACADM* | chr1 | 75733051:75733149 |
| 9.26% | *ZNF638* | chr2 | 71393394:71393645 |
| 6.31% | *PPP1R9A* | chr7 | 95274085:95274168 |
| 6.08% | *SNHG14* | chr15 | 25065122:25065454 |
| 4.11% | ***ACSL6*** | chr5 | 131991745:131991813 |
| 3.89% | *GET1-SH3BGR* | chr21 | 39499823:39499915 |
| 3.82% | *SDR39U1* | chr14 | 24441674:24441795 |
| 3.36% | *ZNF91* | chr19 | 23374638:23374764 |
| 3.00% | *NEBL* | chr10 | 20850395:20850502 |
| 2.52% | ***RIMS2*** | chr8 | 104094478:104094729 |
| 2.34% | ***TULP4*** | chr6 | 158501678:158504178 |
| 2.28% | *STX16-NPEPL1* | chr20 | 58691405:58691562 |
| 2.08% | *ZBTB47-AS1* | chr3 | 42612292:42612367 |
| 1.92% | *ENSG00000268575* | chr1 | 1704223:1704344 |
| 1.60% | *LRRC8C-DT* | chr1 | 89587903:89588085 |
| 1.52% | *NAP1L4* | chr11 | 2989129:2989271 |
| 1.14% | ***KIF1B*** | chr1 | 10303163:10306464 |
| 1.06% | *HECTD1* | chr14 | 31128612:31129448 |
| 0.79% | ***WASF3*** | chr13 | 26676549:26676724 |
| 0.37% | ***PHF14*** | chr7 | 10982372:10983159 |
| 0.35% | *ADD1* | chr4 | 2867972:2868129 |
| 0.30% | *PDE4DIP* | chr1 | 149012591:149012776 |
| 0.16% | *RNF157-AS1* | chr17 | 76153125:76153788 |

**Figure S2. Contribution of variance from brain regions, individuals, and *APOE* genotype to ASAS patterns.**
The percent of variance a brain region (Region), individual, or *APOE* genotype (*APOE*) contributes to for ASAS events for exons identified in at least 2 individuals per brain region and 2 brain regions per individual (Methods). Each row represents an exon, and the columns represent the percent variance of the different categories. Exons are colored by which category the highest percent variance was from. Exons with the highest percent variance from the *APOE* genotype and the percent variance greater than 0.1% are shown in the table. AD-relevant gene names are in bold. Ensembl IDs are shown for genes without approved HGNC symbols. All coordinates are from the hg38 human genome assembly.

### Figure S3

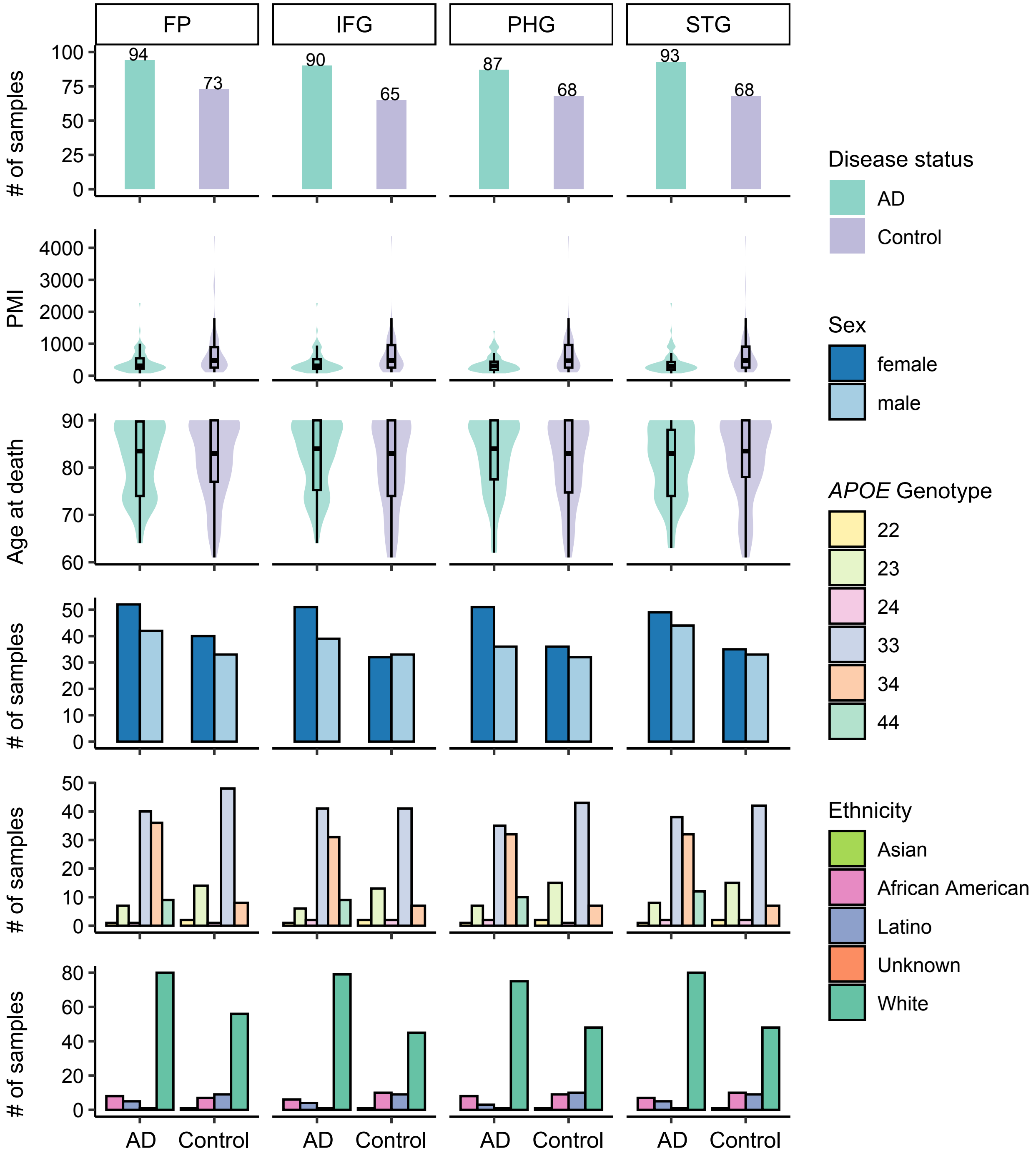


**Figure S3. Distribution of sample size and cohort demographics for identifying putative functional SNPs.**
Sample size, post mortem interval (PMI), age at death, sex, *APOE* genotype, and ethnicity distributions for each brain region in AD and Control groups, based on "definite AD" and "normal" CERAD scores, respectively (Methods). STG: superior temporal gyrus; PHG: parahippocampal gyrus; IFG: inferior frontal gyrus; FP: frontal pole.

### Figure S4

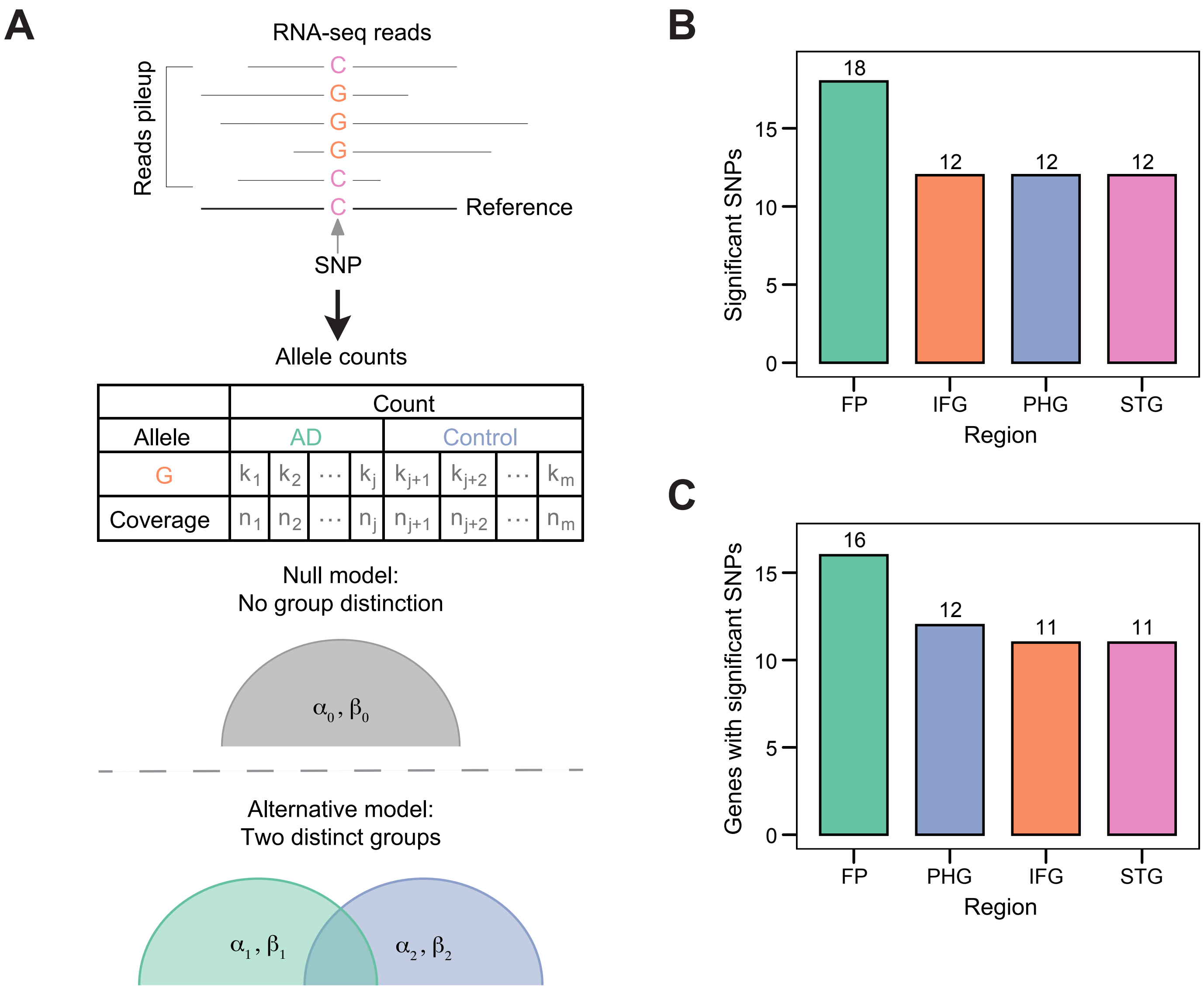


**Figure S4. Identification of tag SNPs displaying differential allelic bias in AD.**
(A) Schematic for identifying differential allelic bias of tag SNPs (Methods). The allelic ratio was modeled as a beta-binomial distribution using a likelihood ratio test to determine significance between AD and control groups. (B) Number of significant differential tag SNPs identified in each region. (C) Number of genes with at least one significant differential tag SNP identified in each region. FP: Frontal Pole; PHG: Parahippocampal Gyrus; STG: Superior Temporal Gyrus; IFG: Inferior Frontal Gyrus.

### Figure S5

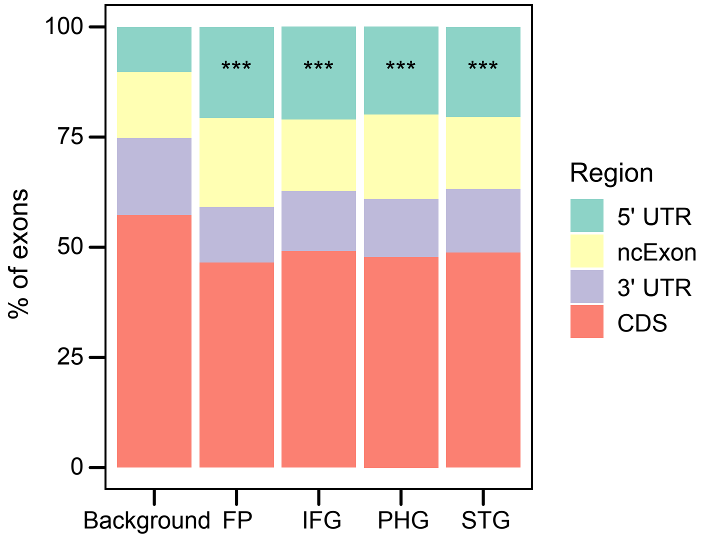


**Figure S5. 5’ UTR enrichment for ASAS exons in each brain region.**
Genomic regions overlapping ASAS exons in each brain region compared to all annotated exons (Background). ***p < 0.001 (Fisher’s exact test). CDS: coding sequence. ncExon: exons in non-coding transcripts. FP: Frontal Pole; PHG: Parahippocampal Gyrus; STG: Superior Temporal Gyrus; IFG: Inferior Frontal Gyrus.

### Figure S6

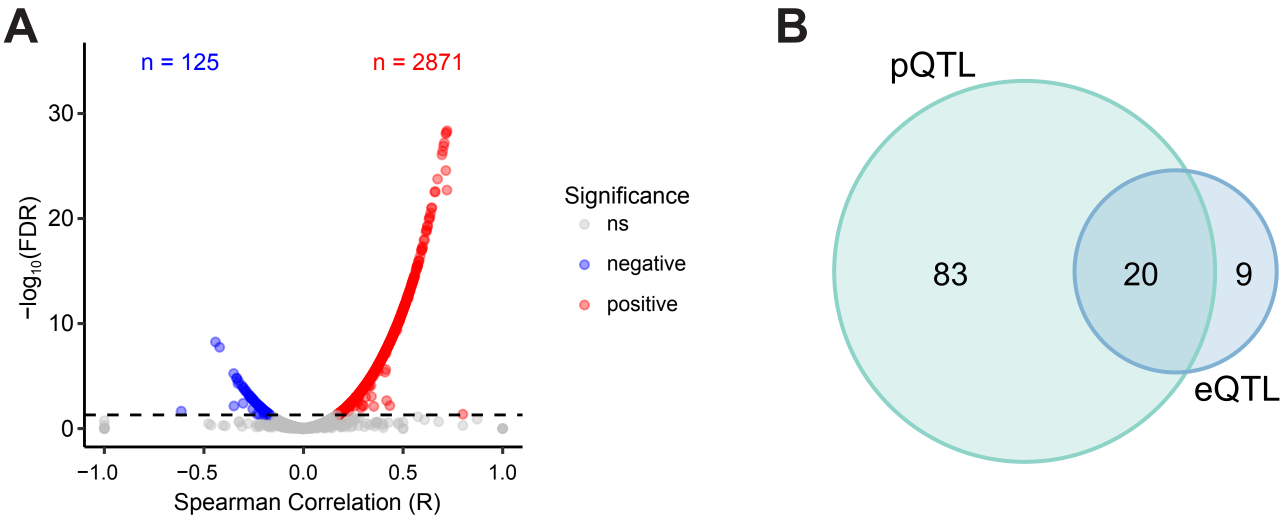


**Figure S6. Relationship between mRNA and protein abundance**
(A) Significance and directionality (Spearman correlation coefficient) of the correlation between protein abundance and RPKM for each gene (Methods). Each dot represents one gene. Red and blue colors represent genes with significant mRNA and protein abundance correlation, in the positive and negative direction, respectively (FDR ≤ 0.05). ns: not significant. (B) Overlap of 5’ UTR ASAS-associated SNPs and pQTLs (Methods) or GTEx brain eQTLs within the same gene.
